# Olive mill wastewater as a source of by-products promoting plant defense against microbial pathogens

**DOI:** 10.1101/2024.06.13.598812

**Authors:** Ascenzo Salvati, Fabio Sciubba, Alessandra Diomaiuti, Gian Paolo Leone, Daniele Pizzichini, Daniela Bellincampi, Daniela Pontiggia

**Affiliations:** Department of Biology and biotechnologies “Charles Darwin”, Sapienza University of Rome, Italy; Department of Environmental Biology, Sapienza University of Rome, Italy; NMR-based Metabolomics Laboratory, Sapienza University of Rome, Italy; Research Center for Applied Sciences for the Protection of the Environment and Cultural Heritage, Sapienza University of Rome; Laboratory Bio-products and Bio-processes BIOAG, ENEA Centro ricerche Casaccia, Italy

**Keywords:** Multi-Phase Decanter, Olive Mill Waste, Pâté’, vegetation water, tangential-flow membrane filtration technology, bioactive molecules-enriched fractions, oligogalacturonides, *Arabidopsis thaliana*, microbial pathogens, plant immunity

## Abstract

Olive oil is a core component of the Mediterranean diet known for its nutritional properties and health benefits. Olive industry is moving to novel extraction systems for higher oil yield and quality and for waste reduction, which is a relevant problem in the process due to its toxicity and high disposal costs. Multi-Phase Decanter (DMF) is a modern two-phase system performed without adding water during the process. Using DMF, a wet by-product indicated as pâté and consisting of the fruit pulp and vegetation water (VW) is recovered. The pâté has a high content of potentially bioactive molecules that may be exploited to promote plant resistance against microbial pathogens. In this work, to identify by/products of biological interest, the VW recovered from the pâté by centrifugation was subjected to fractionation by tangential-flow membrane filtration (TFMF), combining microfiltration (MF) and ultrafiltration (UF). High-resolution NMR spectroscopy indicated the presence of bioactive molecules such as flavonoids, hydroxytyrosol and oleuropein with known antimicrobial activity. High-Performance Anion Exchange Chromatography with Pulsed Amperometric Detection (HPAEC-PAD) was performed to detect the presence of pectic oligosaccharides in the fractions, showing the enrichment, in the UF-concentrate fraction, of oligogalacturonides (OGs), well known for the ability to elicit defense responses and protect plants against pathogen infections. *Arabidopsis thaliana* plants treated with TFMF fractions displayed induction of defense responses and exhibited resistance against microbial pathogens without adverse effects on growth and fitness. This study shows that pâté by-products can potentially be exploited in agriculture as sustainable plant phyto-protectant.

**Graphical abstract:** (image created with BioRender)

## 1. INTRODUCTION

Olive oil is a core component of the Mediterranean diet known for its high nutritional properties and health benefits (Owen *et al*., 2000, Markellos *et al*., 2022). The olive oil industry generates, in a short period of usually four months, huge amounts of olive pomace and olive mill wastewater (OMW), which content vary according to the oil extraction system (Sciubba *et al*., 2020). Olive pomace by- products are easy to recycle and are widely reused both in the food industry, for the production of pomace oil and in the energy industry, for the production of pellets and biomethane. Conversely, OMW, due to its high pollutant charge, is an expensive waste to dispose. OMW is characterized by high acidity (pH of about 5), high levels of biochemical oxygen demand and chemical oxygen demand and contains high concentrations of recalcitrant compounds such as lignin and tannins (52-180 mg/L) (Paraskeva and Diamadopoulos, 2006, Mohammed, 2021). On the other hand, OMW contains a wide range of valuable phenolic compounds with proven antimicrobial properties (e.g. flavonoids, hydroxytyrosol, oleuropein) and OMW by-products are effective as biopesticides against different microbial pathogens (El-Abbassi *et al*., 2017, Russo *et al*., 2022, Jarboui *et al*., 2024). However, the effect of this liquid waste as plant elicitor of defense responses has not yet been investigated.

The increased demand and advances in technology are driving the evolution of olive oil extraction methods with the aim of increasing olive oil yield, ensuring a high quality of the product obtained and reducing amount and disposal costs of OMW (Sciubba *et al*., 2020). In particular a new two-phase decanter, called Multi-Phase Decanter (DMF), has recently been developed and largely adopted in Italian oil mills (Di Giacomo and Romano, 2022). This technology does not require the addition of water during the extraction process and introduces a pulp-kernel separation system producing a kernel-enriched fraction and a high hydrated by-product (moisture content: 75–90%), called pâté, consisting of the fruit pulp and vegetation water (VW). Pâté represents a novel by-product, approx. 45–55% of the total olive weight (Durante *et al*., 2020), rich in water-soluble bioactive molecules that are more concentrated than olive mill by-products obtained with other extraction systems (Lozano-Sánchez *et al*., 2017). The use of Pâté in the food and feed sectors have been recently evaluated (e.g. increasing of food nutritional value and feed fortification) (Foti *et al*., 2022), while the application of Pâté and its by-products in agriculture has not been sufficiently explored, as well as their effects in protecting plants against pathogens are yet not known (Sciubba *et al*., 2020, Benaddi *et al*., 2023). In this work, by tangential-flow membrane filtration (TFMF) the VW, recovered from Pâté, was sequentially fractionated through a combination of microfiltration (MF) and ultrafiltration (UF). In order to evaluate the possible valorization of the VW-derived fractions in plant protection against pathogens we initially exploited analytical platforms to finely characterize the presence of bioactive molecules.

The phytochemical characterization by NMR showed the presence of flavonoids, hydroxytyrosol and oleuropein with a known antimicrobial activity. Our glycomic analysis indicates the presence, in some fractions, of pectin-derived oligogalacturonides (OGs) with a degree of polymerization (DP) between 10 and 17, which are known to act as damage-associated molecular patterns (DAMPs) by eliciting cell wall (CW)-mediated immunity system leading to the activation of defense responses and resistance against pathogens (Osorio *et al*., 2011, Ferrari *et al*., 2013, Pontiggia *et al*., 2020, Gamir *et al*., 2021, De Lorenzo and Cervone, 2022). Noteworthy, externally applied OGs trigger various defense mechanisms that include the buildup of phytoalexins (Davis *et al*., 1986), the activation of defense-related genes (Denoux *et al*., 2008, Osorio *et al*., 2011), the generation of reactive oxygen species (Galletti *et al*., 2008) and jasmonate biosynthesis following cytosolic Ca^+2^ signaling (Moscatiello *et al*., 2006).

We show here the potential of TFMF fractions as elicitors of defense responses in *Arabidopsis thaliana* against the necrotrophic pathogens *Botrytis cinerea* (*Bc*) and *Pectobacterium carotovorum* (*Pc*) after foliar treatment.

## 2. EXPERIMENTAL SECTION

### 2.1. Plant materials and pâté initial processing

*A. thaliana* seeds (ecotype Columbia, Col-0) were purchased from LehleSeeds (Round Rock, TX; USA). Olive drupes (*Olea europaea* L cv. Caninese) were harvested in Northern Lazio (Italy), at Jaén maturation index of 3.44 (Ortenzi *et al*., 2021) and processed (F.lli Palocci S.r.l., Vetralla, Viterbo, Italy) using the Multi-Phase Decanter (DMF) technology (Leopard, Pieralisi, Jesi, Italy).

Pâté (40 kg) was sampled and centrifuged at 5000xg for 20 min. The liquid supernatant (vegetation water, VW) was separated from the pellet, collected and stored at −20°C until the membrane filtration steps were performed.

### 2.2. Tangential-flow membrane filtration (TFMF)

Filtration was conducted on two different bench-scale filtration facilities with a daily treatment capacity of about 5 L/d equipped with a refrigeration loop system to maintain the operating temperature around 20 °C. The microfiltration (MF) unit was an in-house assembled equipment including a peristaltic pump (Masterflex XX80EL230-Millipore Burlington, Masachusetts, USA) and a housing containing a TAMI (TAMI Industries, FR) tubular ceramic membrane with a filtering surface of 0.12 m^2^ and 23 channel structural configuration; the molecular weight cut-off (MWCO) of membrane is 140 nm (about 100 kDa) for the MF trials (Erickson, 2009). Ultrafiltration (UF) was conducted with a flat-sheet membrane-testing module (SEPA CF system, SEPRA S.r.l. Cesano Maderno, MB, Italy) used to obtain performance data on membrane. The lab-scale tangential flow filtration unit, equipped with a volumetric pump with a maximum flow rate of 400 L/h and a maximum pressure 150 bar (Smem, T3A 100 LA4, MB Italy), is made of stainless steel and can accommodate a 137 cm^2^ flat sheet of membrane: a polyethersulfone membrane with a MWCO of 50 kDa (4.8 nm) were used for UF (Erickson, 2009).

Transmembrane pressure (TMP) was calculated through the following equation:

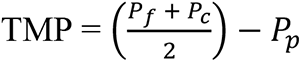

where P_f_ represents the feed stream’s inlet pressure, P_c_ represents the concentrate stream pressure, and P_p_ represents the permeate stream pressure. All measurements are in bars.

During the membrane filtration process, aliquots were taken from the permeate or concentrate TFMF fractions (CMF, PMF, CUF and PUF) and a volume corresponding to 1 g of each fractions was subjected to analysis on a Sartorius MA 160 Infrared moisture analyzer (Sartorius, Göttingen, Germany) at 105°C until reaching 0.1% of residual water. Three replicates of each sample were analyzed. The fractions were lyophilized (Freeze Dryer Edwards) at −40 °C until complete drying, and stored at −20°C. At the end of each filtration phase, permeates and concentrates were collected, aliquoted in 50 ml falcon tubes and stored at –20°C.

Quantity of starting pâté material and obtained pellet, and volumes of the different fractions obtained from the fractionation process, taking into account also the volumes used in the moisture content analysis, are reported respectively in grams and Liters in Fig.1. The overall volume concentration ratio (VCR) for each trial was obtained from the following formula:

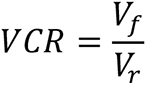

where V_f_ and V_r_ are the volume of the feed and retentate solutions.

**Figure 1.**
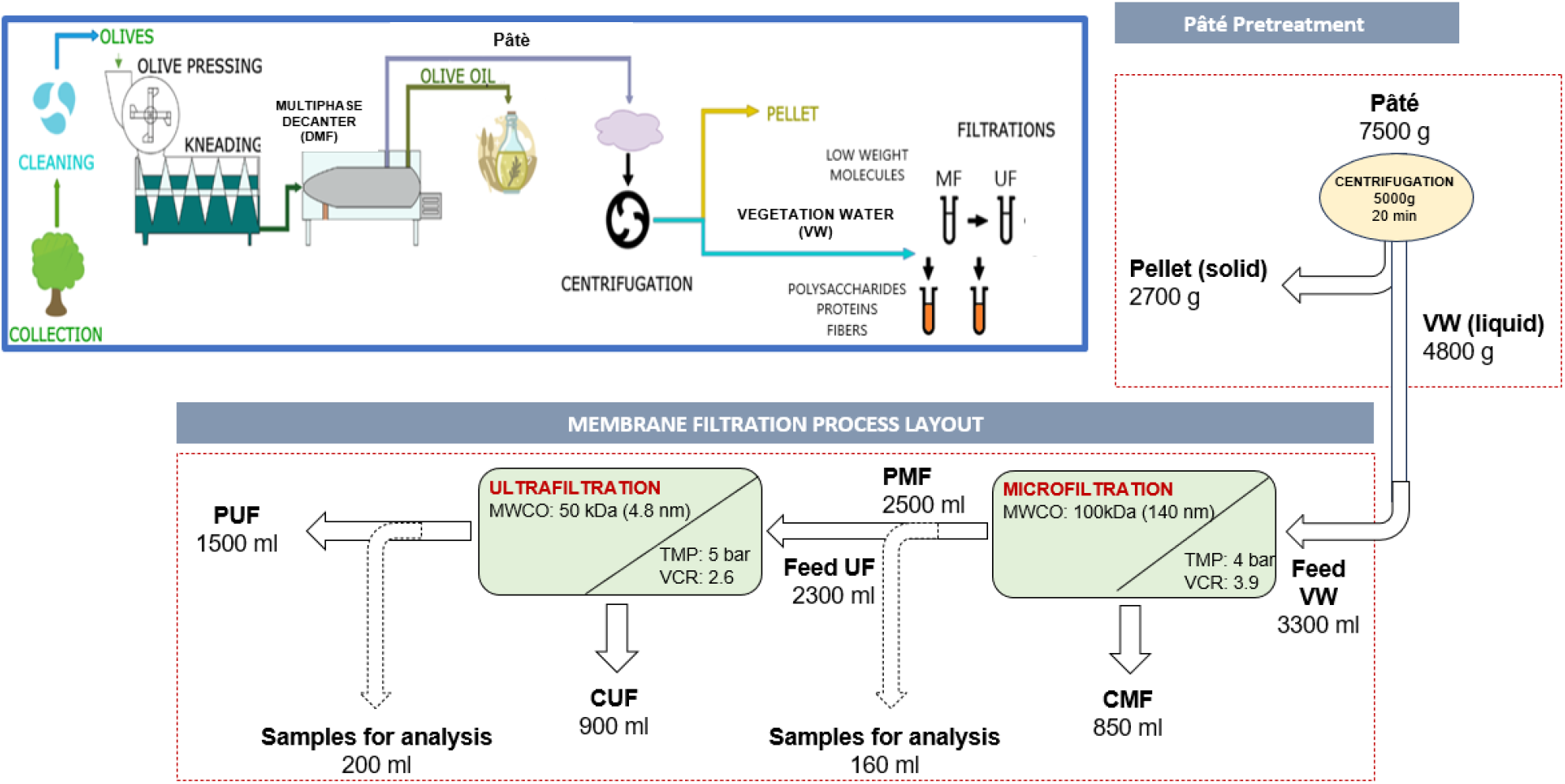
Schematic representation of a single trial of Vegetation Water (VW) fractionation through tangential-flow membrane filtration (TFMF). PMF: permeate of microfiltration, CMF: concentrate of microfiltration, PUF: permeate of ultrafiltration, CUF concentrate of ultrafiltration, MWCO: molecular weight cut-off, Membrane pore diameter is in parentheses, VCR: volume concentration ratio, TMP: transmembrane pressure.

### 2.3. NMR Characterization

Aliquots of VW and of each fraction (CMF, PMF, CUF and PUF) were extracted according to a Blight-Dyer protocol (Giampaoli *et al*., 2021), optimized for the type of sample. Briefly, a mixture of chloroform: methanol: water (1:1:0.6 v/v/v) was added to each sample (1 g). The samples were then vortexed and stored at 4 °C overnight before being centrifuged at 10,000 rpm at 4°C for 20 minutes. After centrifugation, the upper hydroalcoholic fraction and the lower chloroform fraction were separated, brought to dryness under a gaseous nitrogen flow and then stored at −80 °C pending analysis. The samples of the hydroalcoholic fraction were redissolved in 700 µl of D_2_O containing as reference concentration and chemical shift 3-(trimethylsilyl)-2,2,3,3-d 4 -propionate sodium at a concentration of 2 mM. The corresponding chloroform samples were redissolved in 700 µl of CDCl_3_ containing hexamethyldisiloxane as a concentration and chemical shift reference at a concentration of 2 mM.

The spectra were acquired on a JNM-ECZ 600R spectrometer, equipped with a 14.09 Tesla magnet with an operating frequency for hydrogen of 600 MHz and a multinuclear head. The one-dimensional spectra were acquired with a given number of points equal to 64k, a spectral amplitude of 15 ppm (corresponding to 9000 Hz), 128 scans and a recycle time of 7.7 seconds for a total of 15 seconds per scan to guarantee the complete relaxation of the resonances. The two-dimensional TOCSY experiments were acquired with a dot matrix of 8k x 256, a spectral width of 15 ppm in both dimensions, 96 scans, a recycling time of 2 seconds and a mixing time of 80 milliseconds. The two-dimensional HSQC experiments were acquired with a dot matrix of 8k x 256, a spectral amplitude of 15 ppm for hydrogen and 250 ppm for carbon (corresponding to 37500 Hz), 96 scans, a recycling time equal to at 2 seconds and a J ^1^H-^13^C coupling constant of 145 Hz.

The molecule amounts are reported as mg of molecule for g of dry weight of the starting sample.

### 2.4. Extraction of cell wall polysaccharides and analysis of monosaccharide composition

Cell wall polysaccharides isolation from VW and filtrates was performed as previously described with modifications (Lionetti *et al*., 2015). After rehydration with ultrapure water, the freeze-dried samples were homogenized for 1 min at 30 Hz using a mixer mill (MM301; Retsch) and inox beads (5 mm diameter). Milled samples were extracted with pre-warmed (70°C) ethanol 70% (v/v) for 30 min and centrifuged at 14000xg for 10 min. Residues were suspended twice with chloroform: methanol mixture (1:1 v/v) for 30 min and centrifuged as above. Residues were resuspended twice in aqueous solution of acetone 80% (v/v) and centrifuged as above. The final residue, defined alcohol insoluble solid (AIS, total cell walls), was dried using a stream of N_2_ gas at room temperature. Monosaccharide composition of the cell wall matricial component of AIS, mainly composed of pectin and hemicellulose, was determined, after TFA (trifluoroacetic) hydrolysis, by HPAEC-PAD, as previously described (Costantini *et al*., 2023).

### 2.5. Extraction and characterization of OGs

For the isolation of OGs, freeze-dried VW and its TFMF derived fractions (100 mg) were rehydrated in 1 mL of ultra-pure water. Three replicates were analyzed for each sample. For OG precipitation, ethanol was added at a final concentration of 40% (v/v) in 0.05 M sodium acetate, pH 5. After stirring overnight at 4°C, the samples were centrifuged at 25000xg for 20 min at 4°C. Pellets were washed by 70% (v/v) cold ethanol and after one more centrifugation the pellets were placed under a vertical flow chemical hood overnight to completely evaporate the ethanol and resuspended in ultra-pure water for the High-Performance Anion-Exchange Chromatographic with Pulsed Amperometric Detection (HPAEC-PAD) analysis (Pontiggia *et al*., 2015). The OG mixture used as standard had been previously prepared (Bigini *et al*., 2024). For enzymatic degradation of pectic fragments, specific TMFM fractions were treated with commercial fungal pectinase from *Aspergillus aculeatus* (Sigma P2611) that contains mainly pectintranseliminase, polygalacturonase, and pectinesterase at a final concentration of 3.8 units μl^-1^. Samples were sequentially incubated at 30 °C for 3 or 24 h to promote the enzymatic degradation and the enzymatic reaction was then stopped by heating sample at 70°C for 10 min.

### 2.6. Cytosolic free calcium concentration ([Ca^2+^]_cyt_) detection

Ten-day-old seedlings of *A. thaliana* stably expressing the Ca^2+^ biosensor GCaMP3 (35S::GCaMP3) (Grenzi *et al*., 2023)], grown in MS/2 liquid medium, were transferred into black 96-well plate for the analysis. In each well two seedlings were placed in 150 μL of MS/2 liquid medium and 50 μL of freeze-dried VW-derived fractions, resuspended in ultrapure water, were added. Water and solutions containing oligogalacturonides (OGs) were used as negative and positive control, respectively. The [Ca^2+^]_cyt_ detection was performed by GloMax-Multi Detection System (GMDS; Promega Corporation). [Ca^2+^]_cyt_ was detected by measuring the fluorescence emitted at intervals of 1 minute.

### 2.7. Elicitation and gene expression analyses

Ten-day-old *A. thaliana* seedlings, grown in MS/2 liquid medium in 12-well plates, were treated for 1 h with VW, CUF and PUF at the concentration of 300 μg/mL or water as a control. Total RNA was extracted using RNA isolation NucleoZol (Macherey-Nagel) according to the manufacturer’s instructions and treated with RQ1 RNase-free DNase (Promega). cDNA was synthesized with ImProm-II™ Reverse Transcription System (Promega). qRT-PCR was performed with a CFX96 Real-Time PCR System (BioRad) using iTaq Universal SYBR Green Supermix (BioRad) as recommended by the manufacturer. The amplification protocol consisted of 30 s of initial denaturation at 95°C, followed by 45 cycles of 95°C for 15 s, 58°C for 15 s and 72°C for 15 s. The expression levels of each gene, relative to *UBQ5*, were determined using a modification of the Pfaffl method (Pfaffl, 2001) as previously described (Gramegna *et al*., 2016). Sequences for all primers used for quantitative PCR and identifiers of the corresponding genes are listed in Table S1. The data show means of relative expression ± SD (n=3).

### 2.8. Pathogenicity assays

*A. thaliana* plants (Col-0) were grown in a growth chamber at 22 °C, 70% humidity, under irradiance of 100 μE·m^-2^·s^-1^ with a photoperiod of 12-h light/12-h dark. Leaves of 5-week-old *A. thaliana* plants were sprayed with solutions obtained redissolving and subsequently filter-sterilizating freeze-dried VW, CUF and PUF (each at a final concentration of 3 mg/mL). Control plants were sprayed with sterile water. *Botrytis cinerea* and *Pectobacterium carotovorum* subsp. *carotovorum* (strain DSMZ 30169) were inoculated 24 h after treatment. *B. cinerea* conidia were cultured in potato dextrose medium (PDB, 24g/L; Difco, Detroit, USA) at the dilution of 1 × 10^5^ conidia/mL and inoculated on Arabidopsis leaves essentially as described by (Benedetti *et al*., 2015). *P. carotovorum* was grown and inoculated as previously described (Gramegna *et al*., 2016). Symptoms caused by the microbial pathogens were assessed by measuring the area of macerated tissue, at 48 and 16 hpi respectively, using ImageJ software.

### 2.9. Pathogen growth assay

*B. cinerea* growth assay was performed in a multi-well plate containing 2 ml of Potato Dextrose Broth (PDB) medium supplied with 3 mg/ml of the different fractions. Each well was inoculated with 0.5×10^4^ conidia/mL. Four replicates were prepared for each sample. Plates were incubated at 22°C and after 48 h the fungal growth area was measured using ImageJ software.

*P. carotovorum* assay was performed in 50-ml centrifuge tubes containing 5 ml of Luria-Bertani (LB) liquid medium (Duchefa Biochemie, Haarlem, The Netherlands) and the different fractions (at a concentration of 3 mg/ml) or water as a control. Bacterial growth was carried out on a rotary shaker (160 rpm) at 28°C using 1.4*10^7^ cell/mL as starting inoculum and the optical density at 600 nm was measured before incubation and every hour after incubation until the control culture reach the stationary phase. Three replicates were prepared for each sample.

### 2.10. Vegetative plant growth and fitness analysis

*A. thaliana* WT plants (Col-0) were grown in a growth chamber at 22 °C, 70% humidity, under 100 μE·m^-2^·s^-1^ irradiance with a photoperiod of 12 h light/12 h dark and 16 h light/8 h dark for vegetative plant growth and fitness analysis respectively. Four-week-old plants were sprayed with VW, CUF, PUF (at a concentration of 3 mg/mL) or water as control. For the vegetative plant growth analysis, 6-week-old plants were photographed, harvested and the fresh shoot weight measured. For fitness analysis, seeds were collected and their total weight per plant was measured. Seed length and width were measured by MARViN ProLine (MARViTECH GmbH, Wittenburg, Germany).

### 2.11. Statistical analysis

Analyses were performed using Microsoft Excel or the R environment. t-tests or one-way ANOVA followed by Tukey were used to test for statistical significance. Unless otherwise indicated, P < 0.05 was considered to be significant (n.s., not significant; * p < 0.05; ** p < 0.005; *** p < 0.001).

## 3. RESULTS AND DISCUSSION

### 3.1. Tangential-flow membrane filtration (TFMF) of vegetation water isolated from Pâté

The separation and characterization of biological materials is crucial to achieve the understanding of their properties and define their potential biological application and economical valorization. Pâté was subjected to centrifugation to separate the solid portion (fruit pulp) from the liquid portion (vegetation water, VW). After centrifugation, the supernatant/pellet ratio was 1,8:1 (w/w) and the percentage of VW recovered as a supernatant from the Pâté was about 64%.

Tangential-flow membrane filtration (TFMF) is a powerful tool in bioprocessing, allowing the separation and concentration of active biomolecules into specific molecular pools, such as proteins, sugars and secondary metabolites. By using complex mixtures for filtration, TFMF effectively removes impurities while retaining the desired molecules (Pizzichini *et al*., 2005). The efficiency of TFMF primarily stems from the feed stream (VW in this case) flowing tangentially to the membrane surface. This flow pattern enables continuous solute separation throughout the various stages of filtration and minimizes membrane fouling during the process. Noteworthy, respect to other methods such as solvent extraction and chromatography, TFMF has other benefits such as no need for chemicals, easy scaling up, modularity of the process, but TFMF assures the plenty volumetric harvest of both concentrates and permeates and consequently a quantitative recovery of all molecules contained within them (Zagklis *et al*., 2022). This approach provides a solid protocol and repeatable results about extraction and valorization of valuable compounds in olive oil milling by-products.

The filtration process consisted of two sequential stages: microfiltration (MF) and ultrafiltration (UF). In particular, MF was used to remove suspended solids and fibers in VW while UF was employed specifically with the aim of concentrating oligosaccharides present in the mixture. The general process scheme is depicted in Fig. 1 that summarizes the recovery of the fractions and the volume concentration ratio (VCR). At the end of the filtration process, permeate (PMF and PUF) and concentrate (CMF and CUF) of MF and UF were recovered. Table 1 shows the values of total solids for Pâté and derived fractions. The Pâté has a solids content of 19,57%; this percentage increase up to 30,85 % in the pellet after centrifugation, while the supernatant fraction (VW) show a 9,95 % solids content. VW was used as the feed of the membrane separation process.

**Table 1.** Percentage of total solids, expressed as percentage of dry weight to the weight of the initial sample, and pH of hydrated samples. Mean values ±SD are obtained from three replicates (N=3). VW, vegetation water; CMF, microfiltration concentrate; PMF, microfiltration permeate; CUF, Ultrafiltration concentrate; PUF, ultrafiltration permeate.

| | Total solid (%) ±St. Dev. | pH ±St. Dev. |
| --- | --- | --- |
| **Pâte** | 19.57 ±0.346 | 5.08±0.002 |
| **Pellet** | 30.85 ±1.029 | 5.40±0.005 |
| **VW** | 9.95 ±0.098 | 5.10±0.008 |
| **CMF** | 10.83±0.021 | 5.07±0.003 |
| **PMF** | 8.00±0.204 | 5.06±0.003 |
| **CUF** | 9.41±0.058 | 5.07±0.002 |
| **PUF** | 6.63±0.044 | 5.06±0.003 |

CMF has a solids concentration (10,83%) similar to that of VW, suggesting that the suspended solids portion (retained in the MF) had been efficiently reduced by the centrifugation. PMF and CUF show a concentration of solids of 8% and 9,41% respectively, while in the PUF the amount of solids is 6,63%, about 30% less than CUF. The pH values were acidic and did not significantly vary across all the TFMF process stages. The TFMF approach, here applied at laboratory scale, can be easily scaled-up and it is suitable to be applied also at industrial level (Pizzichini *et al*., 2009, Germani *et al*., 2012).

### 3.2. Phytochemical characterization of VW and TFMF-derived fractions

The biochemical composition of OMWs is highly variable and a fine characterization is needed to define their appropriate valorization (Sciubba *et al*., 2020, Benaddi *et al*., 2023). The phytochemical characterization of VW and the fractions obtained from filtration was carried out employing NMR Spectroscopy. The identification of the molecules in the samples (matrices) was carried out on the basis of 2D experiments as well as literature data (Spinelli *et al*., 2022) and reported in Table S2. A total of 32 molecules belonging to the classes of Amino Acids, Organic Acids, Carbohydrates, Lipids and Secondary Metabolites were identified and 25 were quantified. Only the molecules whose resonances were not superimposed with other signals were chosen for quantification and the results are reported in Table 2.

**Table 2.** Phytochemical composition of vegetation water and its derived fractions after tangential membrane filtration assessed by NMR spectroscopy. Mean values are obtained from three replicates (N=3) with ANOVA significance assessed by Tuckey test. VW, vegetation water; CMF, microfiltration concentrate; PMF, microfiltration permeate; CUF, Ultrafiltration concentrate; PUF, ultrafiltration permeate.

| Molecule | Amount (mg/g DW) |  |  |  |  |
| --- | --- | --- | --- | --- | --- |
|  | VW | CMF | PMF | CUF | PUF |
| Glutamine | 32.34 ± 1.66 a | 54.73 ± 1.67 b | 52.59 ± 1.58 b | 80.44 ± 2.44 c | 48.59 ± 1.61 b |
| Citric acid | 84.56 ± 4.19 a | 81.57 ± 2.46 a | 99.48 ± 3.04 a | 86.94 ± 2.64 a | 76.69 ± 2.34 a |
| Quinic acid | 43.94 ± 2.24 a | 40.39 ± 1.23 a | 53.32 ± 1.58 b | 47.49 ± 1.42 ab | 54.88 ± 1.61 b |
| Malic acid | 61.18 ± 3.02 a | 57.55 ± 2.11 a | 71.3 ± 2.19 b | 60.5 ± 1.83 a | 68.49 ± 2.05 a |
| p-Coumaric acid | 4.97 ± 0.29 a | 5.72 ± 0.18 b | 5.71 ± 0.12 b | 7.83 ± 0.2 c | 3.66 ± 0.15 d |
| Glucose | 265.66 ± 13.25 a | 253.52 ± 7.57 a | 331.11 ± 9.96 b | 285.44 ± 8.54 a | 368.07 ± 10.98 b |
| Sucrose | 9.16 ± 0.49 a | 9.94 ± 0.26 a | 10.32 ± 0.36 a | 14.54 ± 0.41 b | 6.73 ± 0.15 c |
| Stearic acid | 7.7 ± 0.39 a | 25.08 ± 0.79 b | ND | ND | ND |
| Oleic acid | 6.92 ± 0.39 a | 60.45 ± 1.85 b | ND | ND | ND |
| Linoleic acid | 1.66 ± 0.1 a | 24.38 ± 0.7 b | ND | ND | ND |
| Linolenic acid | 0.29 ± 0.1 a | 7.13 ± 0.18 b | ND | ND | ND |
| Triterpenes | 1.17 ± 0.1 a | 16.98 ± 0.53 b | 8.75 ± 0.24 c | 9.76 ± 0.31 c | 25.76 ± 0.73 d |
| Triglycerides | 0.88 ± 0.19 a | 18.04 ± 0.53 b | 4.13 ± 0.12 c | 4.37 ± 0.1 c | 12.44 ± 0.44 d |
| Isopropanol | 1.17 ± 0.1 a | 1.14 ± 0.09 a | 1.34 ± 0.12 a | 1.32 ± 0.1 a | 1.17 ± 0.15 a |
| Choline | 1.36 ± 0.1 a | 1.32 ± 0.09 a | 1.34 ± 0.12 a | 1.53 ± 0.1 a | 1.9 ± 0.15 a |
| Hydroxytyrosol | 16.66 ± 0.88 a | 12.76 ± 0.35 b | 16.52 ± 0.49 a | 18.1 ± 0.51 c | 14.93 ± 0.44 ab |
| Tyrosol | 1.66 ± 0.1 a | 0.35 ± 0.09 b | 0.36 ± 0.12 b | 0.51 ± 0.1 b | 0.15 ± 0.15 c |
| Oleuropein | 128.49 ± 6.43 a | 135.08 ± 4.05 a | 161.79 ± 5.34 b | 162.6 ± 4.88 b | 148.25 ± 4.98 ab |
| Lingstroside | 14.03 ± 0.68 a | 15.49 ± 0.44 a | 17.98 ± 0.49 b | 21.05 ± 0.61 c | 16.68 ± 0.44 b |
| Lingstroside dialdehyde | 9.06 ± 0.49 a | 11.97 ± 0.35 b | 12.51 ± 0.36 b | 11.69 ± 0.31 b | 13.46 ± 0.44 b |
| Oleuropein dialdehyde | 61.18 ± 3.02 a | 69.69 ± 2.11 a | 79.44 ± 2.79 b | 72.4 ± 2.14 b | 84.59 ± 2.49 c |
| Flavonoids | 109.99 ± 5.46 a | 27.37 ± 0.79 b | 34.62 ± 1.09 c | 60.61 ± 1.83 d | 66.3 ± 2.05 d |
| Oleuropien aglycone | 4.38 ± 0.19 a | 4.49 ± 0.18 a | 6.19 ± 0.24 b | 6.81 ± 0.2 b | 18.29 ± 0.59 c |
| Lingstroside aglycone | 0.68 ± 0.1 a | 0.62 ± 0.09 a | 1.09 ± 0.12 b | 1.22 ± 0.1 b | 3.07 ± 0.15 c |
| Pinoresinol | 1.75 ± 0.11 a | 1.67 ± 0.09 a | 2.55 ± 0.12 b | 2.75 ± 0.1 b | 8.34 ± 0.29 c |
| Carotenoids | 5.36 ± 0.19 a | 4.93 ± 0.18 a | 10.08 ± 0.24 b | 10.37 ± 0.31 b | 31.17 ± 0.88 c |

Comparison molecules in the VW fractions showed, as expected, a significant difference between the concentrated fractions (CMF and CUF) and the permeate ones (PMF and PUF), and this is of remarkable importance in terms of bioactive compounds observed: p-Coumaric acid (Lou *et al*., 2012), Tyrosol and Hydroxytyrosol (Bisignano *et al*., 1999), Oleuropein, Lingstroside and their derivatives (Bisignano *et al*., 1999, Medina *et al*., 2006, Macedo *et al*., 2018) and flavonoids (Miho *et al*., 2024). In greater detail, regarding the concentration of these molecules, it was possible to observe that the highest amount of bioactive molecules is observed in the CMF fraction. However, the use of this fraction is undermined by the presence of fatty acids that could negatively affect the performance of products derived from it. In particular, the application of these molecules on soil or leaves could coat them and impede the gas exchanges. Of the other fractions, CUF presents the highest levels of bioactive compounds (Table 2), mainly p-coumaric acid, secoiridoids (oleuropein, lingstroside and their derivatives), hydroxytyrosol and flavonoids, at concentration ranges described to induce biological effects in plants (Luque de Castro and Japón-Luján, 2006, Vagelas *et al*., 2009, Yangui *et al*., 2010, Sciubba *et al*., 2020, Selim *et al*., 2022) therefore suggesting this fraction to be the most promising to test its potential elicitor activity in Arabidopsis.

### 3.3. Monosaccharide composition of AIS in VW and TFMF-derived fractions

The monosaccharide composition of the Alcohol-Insoluble Solids (AIS) of VW and TFMF-derived fractions was determined (Table 3). Monosaccharide composition of VW indicated the presence of pectin-related sugars (arabinose, galacturonic acid, and rhamnose) and hemicellulose-related monosaccharides (including xylose, mannose, and galactose) thus suggesting a relative abundance of Arabinogalactans and Xyloglucans. A large quantity of these polysaccharides seems to be retained in concentrated fractions CMF and CUF. Specifically in CUF, pectins and hemicelluloses possibly, in forms of oligosaccharide fragments, are enriched respect to the permeates making this fraction, particularly interesting for inducing possible biological effects.

**Table 3.**
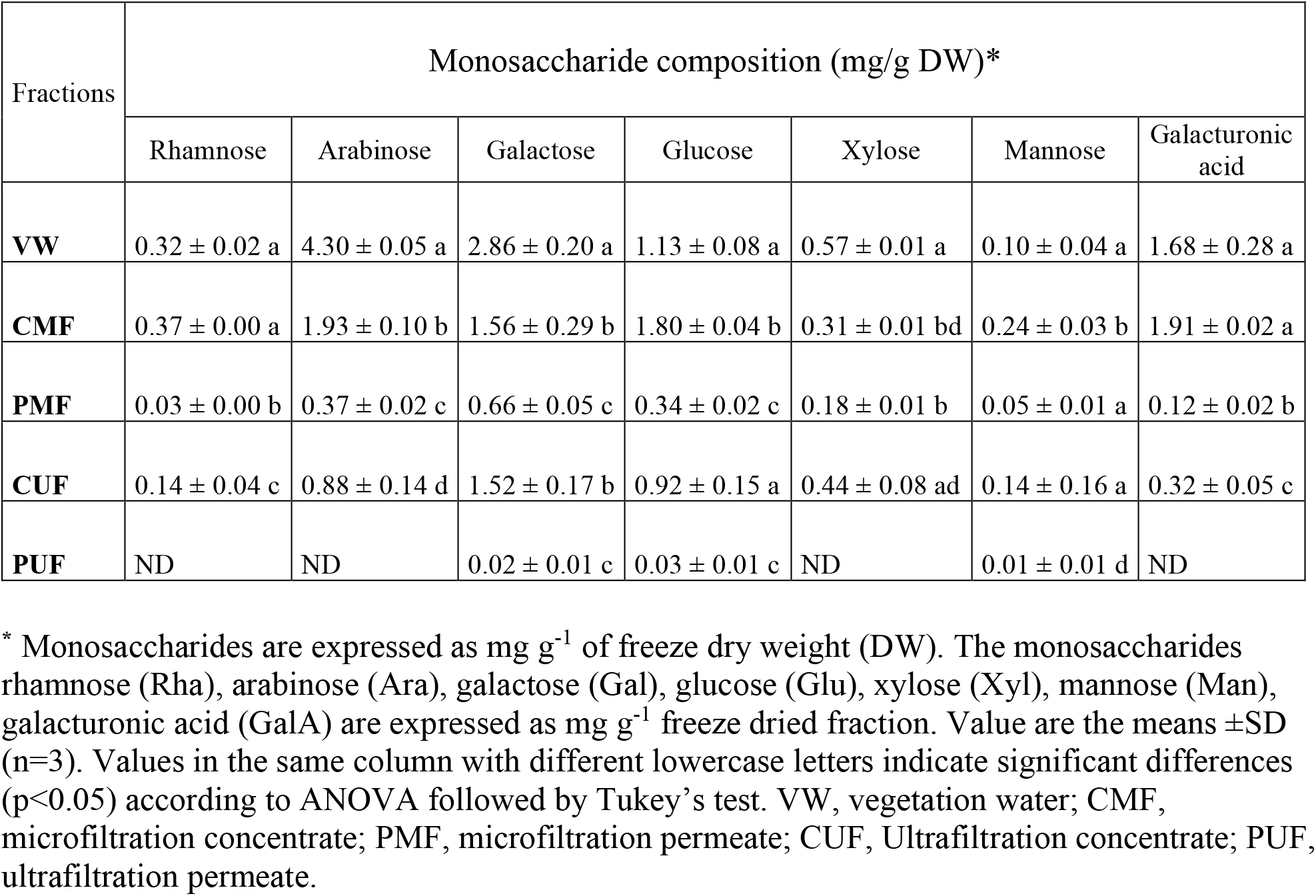
Monosaccharide recovery and monosaccharide composition of the cell wall matricial component of the alcohol insoluble solids (AIS) from vegetation water and its derived fractions after tangential membrane filtration.

| Fractions | Monosaccharide composition (mg/g DW)* |  |  |  |  |  |  |
| --- | --- | --- | --- | --- | --- | --- | --- |
|  | Rhamnose | Arabinose | Galactose | Glucose | Xylose | Mannose | Galacturonic acid |
| **VW** | 0.32 ± 0.02 a | 4.30 ± 0.05 a | 2.86 ± 0.20 a | 1.13 ± 0.08 a | 0.57 ± 0.01 a | 0.10 ± 0.04 a | 1.68 ± 0.28 a |
| **CMF** | 0.37 ± 0.00 a | 1.93 ± 0.10 b | 1.56 ± 0.29 b | 1.80 ± 0.04 b | 0.31 ± 0.01 bd | 0.24 ± 0.03 b | 1.91 ± 0.02 a |
| **PMF** | 0.03 ± 0.00 b | 0.37 ± 0.02 c | 0.66 ± 0.05 c | 0.34 ± 0.02 c | 0.18 ± 0.01 b | 0.05 ± 0.01 a | 0.12 ± 0.02 b |
| **CUF** | 0.14 ± 0.04 c | 0.88 ± 0.14 d | 1.52 ± 0.17 b | 0.92 ± 0.15 a | 0.44 ± 0.08 ad | 0.14 ± 0.16 a | 0.32 ± 0.05 c |
| **PUF** | ND | ND | 0.02 ± 0.01 c | 0.03 ± 0.01 c | ND | 0.01 ± 0.01 d | ND |

\* Monosaccharides are expressed as mg g<sup>-1</sup> of freeze dry weight (DW). The monosaccharides rhamnose (Rha), arabinose (Ara), galactose (Gal), glucose (Glu), xylose (Xyl), mannose (Man), galacturonic acid (GalA) are expressed as mg g<sup>-1</sup> freeze dried fraction. Value are the means ±SD (n=3). Values in the same column with different lowercase letters indicate significant differences (p<0.05) according to ANOVA followed by Tukey's test. VW, vegetation water; CMF, microfiltration concentrate; PMF, microfiltration permeate; CUF, Ultrafiltration concentrate; PUF, ultrafiltration permeate.

### 3.4. PMF and CUF fractions contain biologically active OGs

Many studies indicated that the foliar application of OGs with DP between 10 and 17 protect plants against pathogen infection (Aziz *et al*., 2004, Ferrari *et al*., 2007, Rakoczy-Lelek *et al*., 2023, Bigini *et al*., 2024, Radkowski *et al*., 2024). OGs are well known danger signals that trigger the CW-mediated immunity system leading to the elicitation of defense responses and disease resistance (Ferrari *et al*., 2013, Pontiggia *et al*., 2020).

In order to valorize VW and TFMF-derived fractions as phyto-protectants, whether they contain bioactive OGs was examined by HPAEC-PAD glycomic analysis. For VW and the CMF fraction, chromatographic profiles did not show peaks corresponding to OGs when compared to the chromatographic profile of previously purified OGs mixture enriched in bioactive fragments (Fig. S1), probably due to a very low concentration of these CW fragments or to their possible interaction with other macromolecules (Sieminska-Kuczer *et al*., 2022). Instead, the chromatographic profiles of the PMF and CUF fractions showed, in addition to large oligosaccharides with retention time of more than 26 minutes, an enrichment in oligosaccharides with DP ranging from 10 to 17 units that could be assigned to OGs (Fig. 2, Table S3).

**Figure 2.**
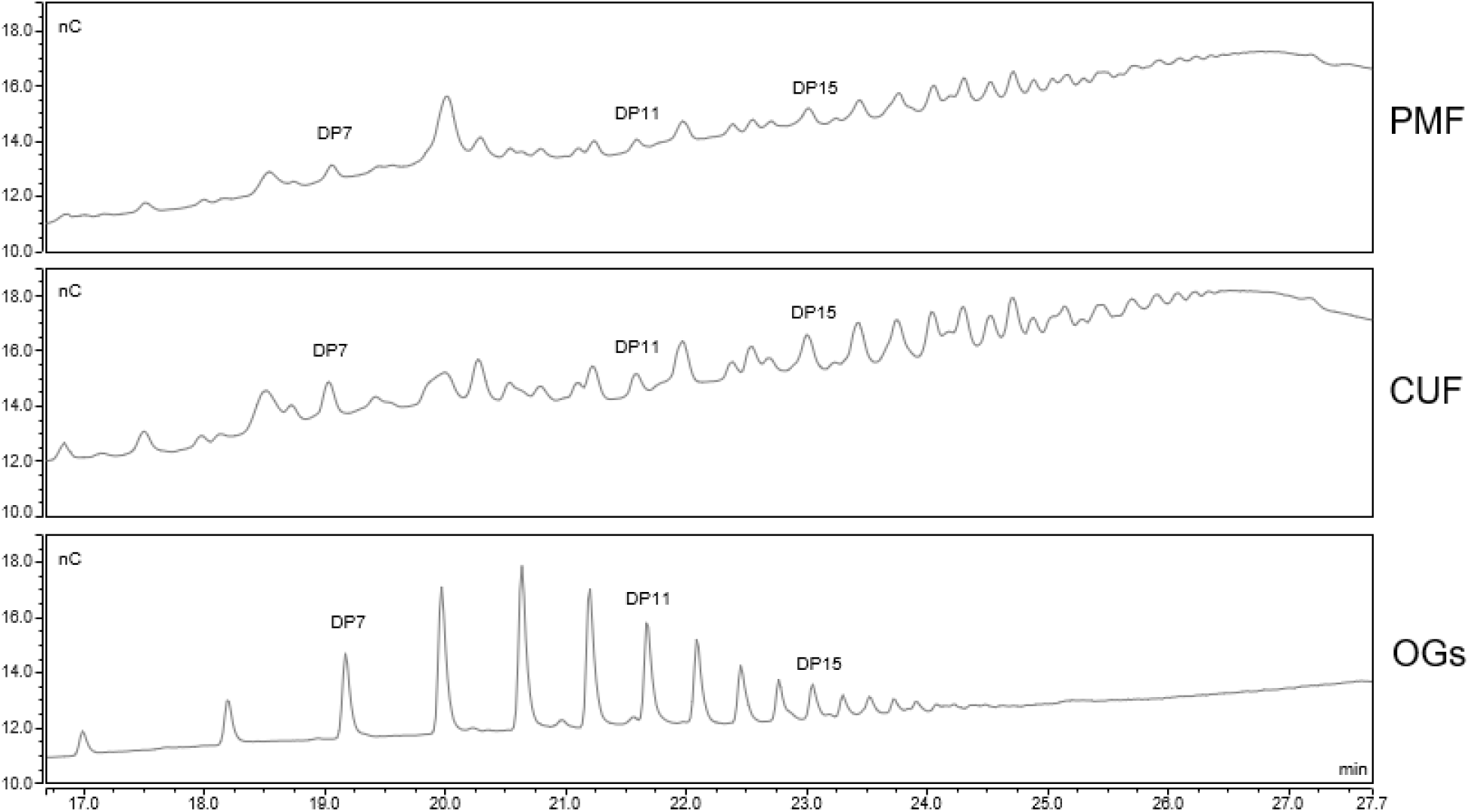
The oligosaccharide profiles of PMF and CUF show peaks corresponding to active OGs. HPAEC-PAD analysis of PMF and CUF fractions compared with purified OGs mixture. Standard OGs are also shown with degree of polymerization (DP). Signal intensities (nC) at each retention time (min) are indicated.

In the PUF fraction, no peaks associated with OG oligomers were evident, indicating that these oligosaccharides were completely retained by the ultrafiltration membrane (Fig.S1).

The nature of the oligomers of interest was assessed by performing a digestion of CUF with a fungal pectinase. The specific hydrolytic enzyme was able to completely degrade the oligosaccharides contained in the sample (Fig. 3), indicating the pectic nature of the oligomers.

**Figure 3.**
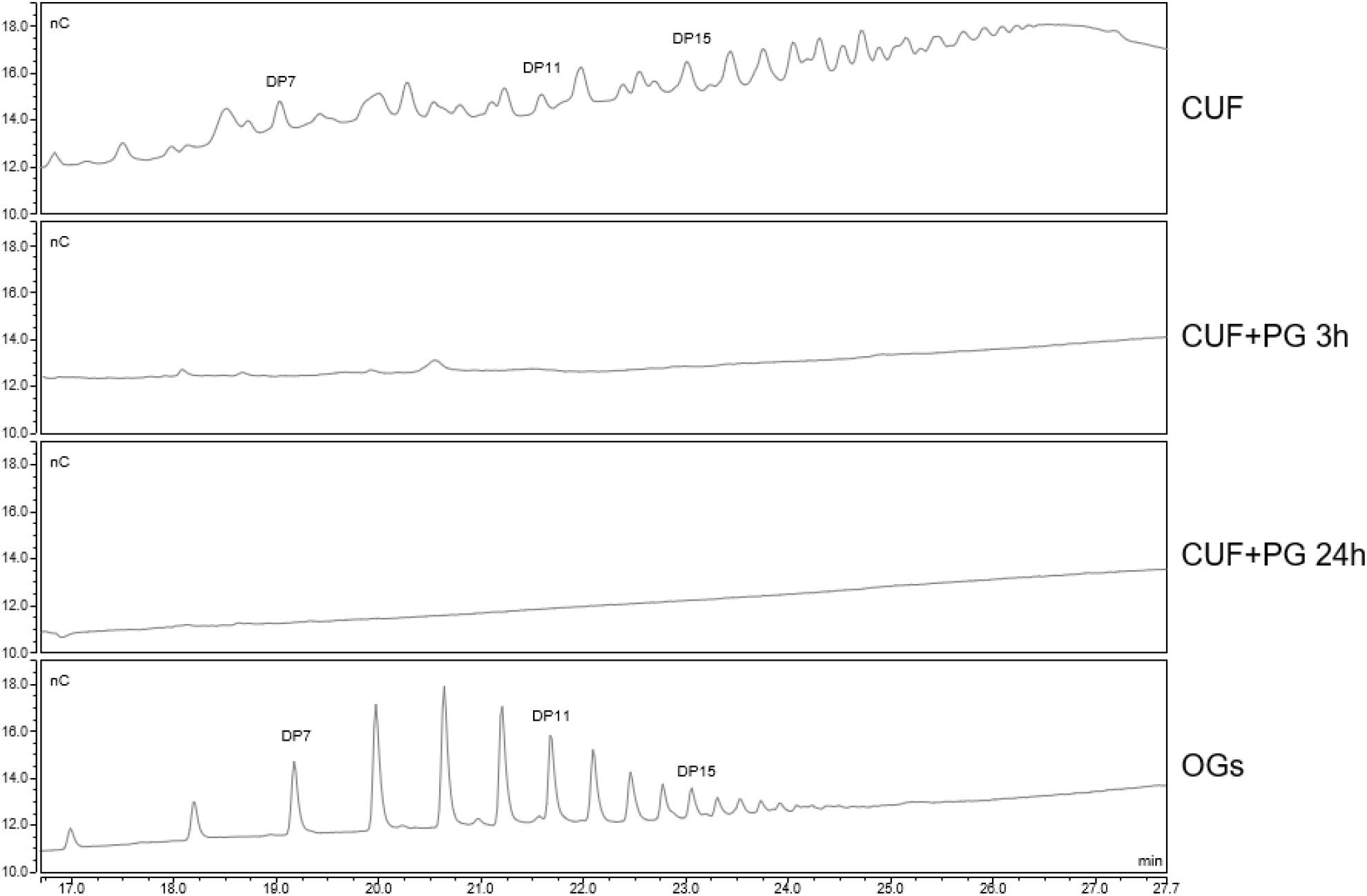
CUF fraction treated with polygalacturonase doesn’t show peaks corresponding to active OGs. HPAEC-PAD analysis of CUF fraction in the absence or presence of fungal pectinase for 3 and 24 hours respectively. Standard OGs are also shown with degree of polymerization (DP). Signal intensities (nC) at each retention time (min) are indicated.

### 3.5. The CUF fraction induces cytosolic free Ca^2+^ ([Ca^2+^]_cyt_) transient increase in *A. thaliana* seedlings

Given the presence of bioactive OGs, we next examined VW-derived TFMF fractions for the capability to induce defense responses in the model plant *A. thaliana*. OGs trigger a wide range of defense responses, including a transient increase in ([Ca^2+^]_cyt_) (Navazio *et al*., 2002, Aziz *et al*., 2004, Moscatiello *et al*., 2006, Wang and Luan, 2024). Calcium ion is firmly established as an ubiquitous intracellular second messenger with a key role in numerous plant defense signaling pathways (Lecourieux *et al*., 2006, Wang and Luan, 2024). Transient increases in [Ca^2+^]_cyt_ can be detected by a wide range of Ca^2+^-binding protein sensors (Batistic and Kudla, 2012, Vigani and Costa, 2019, Grenzi *et al*., 2023).

We first treated transgenic Arabidopsis seedlings expressing the Ca^2+^ biosensor GCaMP3 (Grenzi *et al*., 2023) with active OGs (DP 10-17; 10 and 30 µg/ml) and a rapid single transient of [Ca^2+^]_cyt_ increase was observed within 5 minutes, with a dose-response relationship (Fig. 4A).

**Figure 4.**
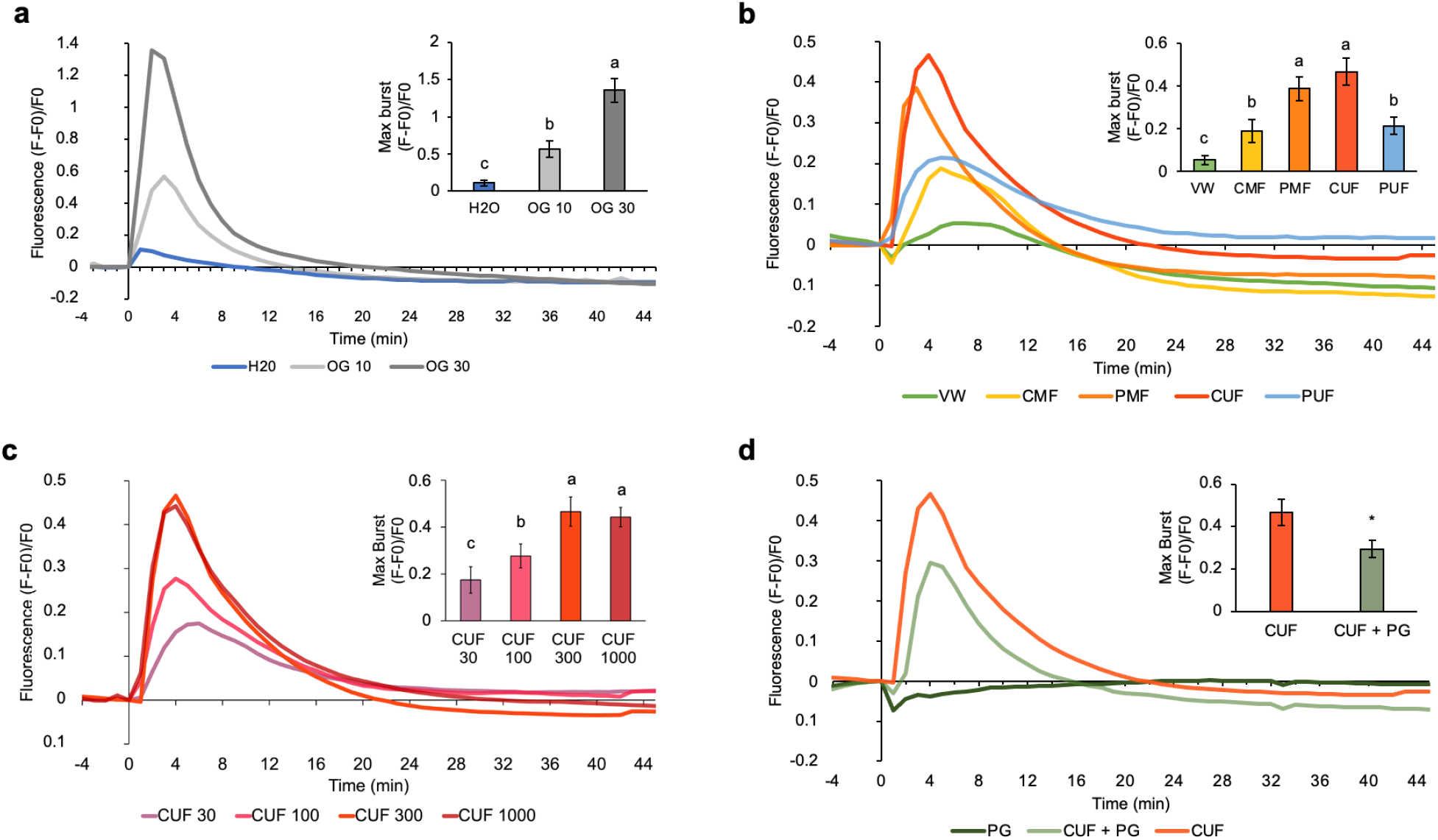
CUF induces cytosolic Ca^2+^ concentration ([Ca^2+^]_cyt_) transient increase similar to that induced by OGs in *A. thaliana*. Representative traces (n=12) of normalized GFP fluorescence (ΔF/F0) in 35S::GCaMP3 seedlings are shown. Traces display 4 min before treatment until 45 min post-treatment. Bars show the maximal [Ca^2+^]_cyt_ values (ΔF/F0) ± SE. OG and CUF concentrations are expressed in μg/mL. (a) OGs trigger concentration-dependent transient increase of [Ca^2+^]_cyt_. (b) CUF triggers the higher [Ca^2+^]_cyt_ transient increase among TFMF fractions. All fractions were used at a concentration of 300 μg/mL. (c) CUF induced [Ca^2+^]_cyt_ transient increase is concentration-dependent. (d) PG-treated CUF at 300 μg/mL induce a significantly lower [Ca^2+^]_cyt_ transient increase than CUF at the same concentration. The little increase of fluorescence induced by H_2_O, shown in figure a, was subtracted to each trace in the figures b, c and d. Asterisks indicate statistically significant differences in data according Student’s t-test (n.s., not significant; * p < 0.05; *** p < 0.001). Different letters indicate P<0.05, as determined by ANOVA with Tukey’s HSD test.

The treatment with VW, as well as with all the TFMF fractions induced [Ca^2+^]_cyt_ transients although at significantly different levels and with different kinetics (Fig. 4B). The higher effect was observed with PMF and was comparable to the effect of the CUF fraction. The PUF fraction, although it does not show OG-like chromatographic peaks, also triggered a significant transient increase in [Ca^2+^]_cyt_, albeit lower than that induced by CUF and PMF and with a different kinetic (Fig. 4B), likely due to presence of other bioactive metabolites (see table 2).

Next, we treated the 35S::GCaMP3 seedlings with the CUF fraction at different concentrations, resulting a dose-dependent increase in [Ca^2+^]_cyt_ with kinetics similar to the OG-induced burst (Fig. 4C). The concentration of 300 μg/mL induced the highest burst, whereas treatment with the highest concentration of 1000 μg/mL did not lead to further increase in fluorescence. Although CUF is a OG-enriched fraction, other bioactive molecules such as phenols are present in quantities that could be also capable of inducing appreciable [Ca^2+^]_cyt_ transients. Indeed, changes in [Ca^2+^]_cyt_ have been reported as ubiquitous event in response to a large array of stimuli (Lecourieux *et al*., 2006, Negi *et al*., 2023). In order to discriminate the effect of OGs from that of other bioactive molecules present in CUF fraction, 35S::GCaMP3 seedlings were treated with CUF digested with pectinase, to specifically degrade the pectic fragments. A significant lower [Ca^2+^]_cyt_ transient increase was observed in seedlings treated with CUF in addition with the pectic enzyme respect CUF alone indicating that OGs contained in the fraction contribute to the production of the [Ca^2+^]_cyt_ signal, although other bioactive molecules present in the fractions appear also to exert this effect (Fig. 4D).

### 3.6. CUF fraction induces defense-related gene expression in *A. thaliana* seedlings

Ca^2+^ signaling induced by elicitors, including OGs, is decoded in plants by an array of Ca^2+^-binding proteins giving rise to a cascade of downstream effects, including transcriptional reprogramming for the implementation of plant defense (Aziz *et al*., 2004, Moscatiello *et al*., 2006, Pontiggia *et al*., 2024, Wang and Luan, 2024). WRKY DNA-binding protein 40 (WRKY40), encoding a transcription factor that acts as a regulator of basal defense (Xu *et al*., 2006), CYP81F2, encoding a cytochrome P450 involved in indol-3-yl-methyl glucosinolate catabolism (Clay *et al*., 2009) and FLG22-INDUCED RECEPTOR-LIKE KINASE 1 (FRK1), the expression of which is considered to be Ca^2+^-independent (Boudsocq *et al*., 2010), have been well characterized as markers of the OG-induced early defense response in Arabidopsis (Denoux *et al*., 2008, Galletti *et al*., 2008, Savatin *et al*., 2014). We analyzed the expression of these genes in seedlings treated for 1 h with CUF and H_2_O, VW and PUF as controls (Fig. 5). As expected, the OG-containing CUF fraction induced the expression of the maker defense genes analyzed. Interestingly, also treatment with VW and PUF induced the expression of marker genes of the early defense response, however at a lower level. These results suggest that, in the CUF fraction, OGs allow stronger activation of plant immunity, working in cooperation with other bioactive compounds present in olive VW derived fractions, possibly phenols and flavonoids.

**Figure 5.**
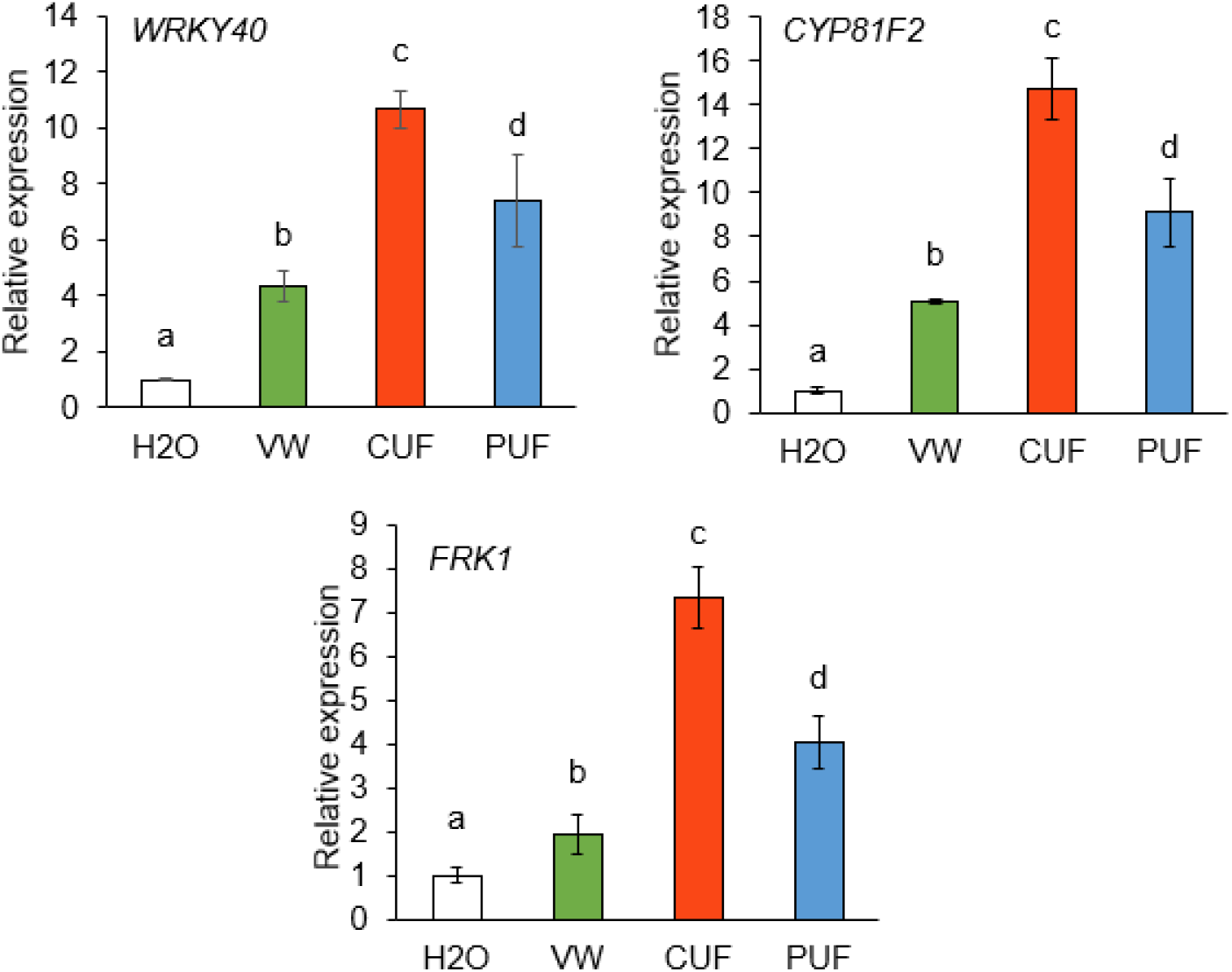
CUF induces high-level expression of defense marker genes in *A. thaliana*. Ten-days-old WT seedlings were treated for 1h with water, VW, CUF and PUF at the concentration of 300 μg ml−1. *WRKY40*, *CYP81F2* and *FRK1* expression was analyzed by qRT-PCR using UBQ5 as reference gene. The data show mean ± SD (n=3) and different letters indicate P<0.05, as determined by ANOVA with Tukey’s HSD test.

### 3.7. Pre-treatment of *A. thaliana* plants with VW, CUF and PUF induces resistance against both fungal and bacterial pathogens

As DAMPs, OGs can induce also various changes at a the physiological, molecular, and epigenetic level which prepare the plant to respond to future stresses. This mechanism is called priming and can be considered an "immunological memory" that allows the plant to be less susceptible to biotic and abiotic stresses as a result of previous stress situations (Martinez-Medina *et al*., 2016, Gamir *et al*., 2021, Zhu *et al*., 2024). *A. thaliana* adult plants pre-treated with OGs show a significant higher resistance to pathogen attack than untreated plants: this induced resistance is evident at 24 hours post-infection and even later (Ferrari *et al*., 2007).

To further evaluate the potential effect of OG-enriched CUF as phyto-protectant against microbial pathogens, WT Arabidopsis plants were sprayed with VW, CUF, and PUF, and 24 hours later, inoculated with *Bc* and *Pc*. Lesion areas were evaluated 48 and 16 hours after infection, respectively (Fig. 6). Both VW and TFMF fractions induced a significant reduced susceptibility against both pathogens compared to the control. While, in CUF fraction, bioactive OGs may contribute to the protective effect against pathogen infection, in PUF the protective effect of the fraction could be attributed to other bioactive molecules, such as phenols and flavonoids. The antimicrobial activity of phenols, has been explored by several authors and has been associated to plant resistance to phytopathogens, suggesting a possible application of these compounds as biopesticides in agriculture (Yakhlef *et al*., 2018, Abbattista *et al*., 2021, Košćak *et al*., 2023, Zhu *et al*., 2024).

**Figure 6.**
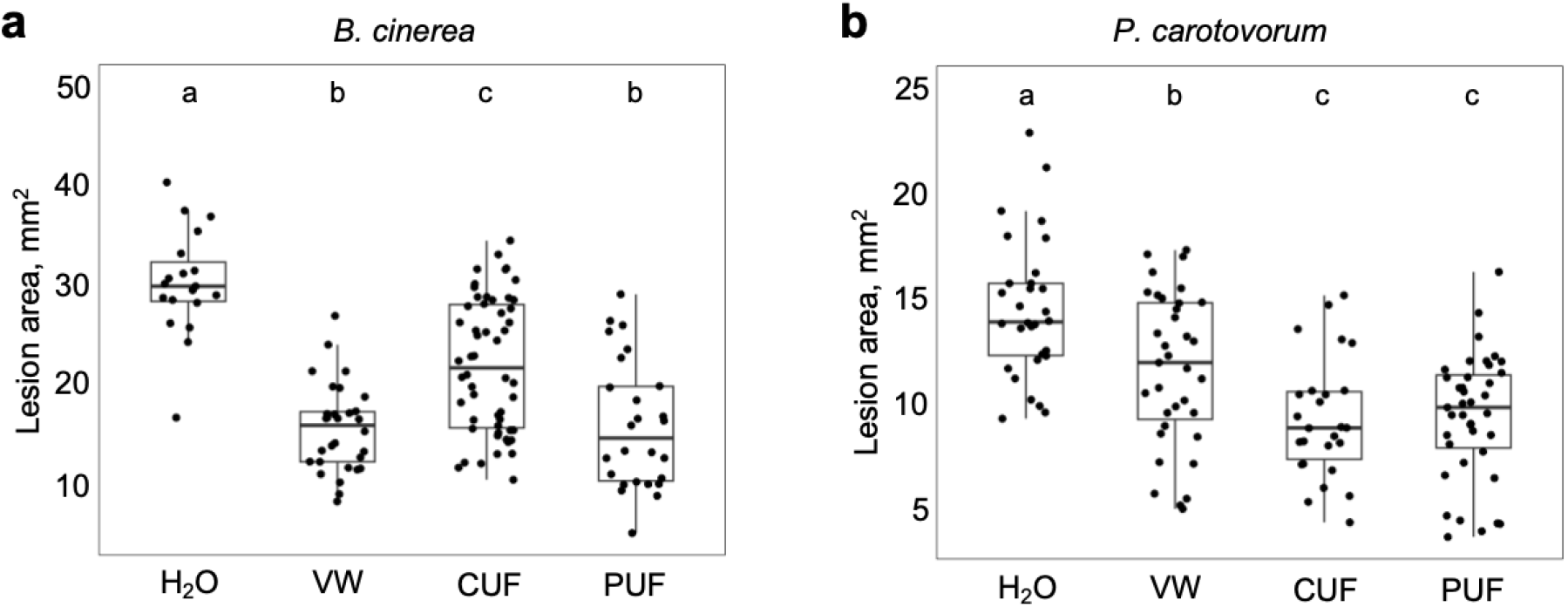
*A. thaliana* plants sprayed with VW, CUF and PUF show resistance to *B. cinerea and P. carovororum*. Freeze-dried VW, CUF and PUF, resuspended in H2O and sterilized, were sprayed 24 h before infection at a final concentration of 3 mg/mL. H2O was sprayed as control. Lesion area produced by *B. cinerea* (a) and *P. carotovorum* (b) was measured, respectively, at 16 and 48 h post infection. The data show mean ± SD (n=3). Different letters indicate P<0.05, as determined by ANOVA with Tukey’s HSD test.

We also tested the direct antimicrobial effect of VW and VW-derived fractions, by evaluating the growth of *Bc* and *Pc* on culture media supplied with VW and its derived fractions. Neither VW nor the derived fractions inhibited *Bc* growth (Fig. 7A), suggesting that the resistance to this fungus in Arabidopsis plants treated with VW and the TFMF fractions is not due to direct antifungal activity. On the other hand, a significant antibiotic activity was observed against *Pc* (Fig. 7B), suggesting that an antimicrobial effect of VW and the TFMF fractions can participate in the induced resistance against the bacterial pathogen in Arabidopsis.

**Figure 7.**
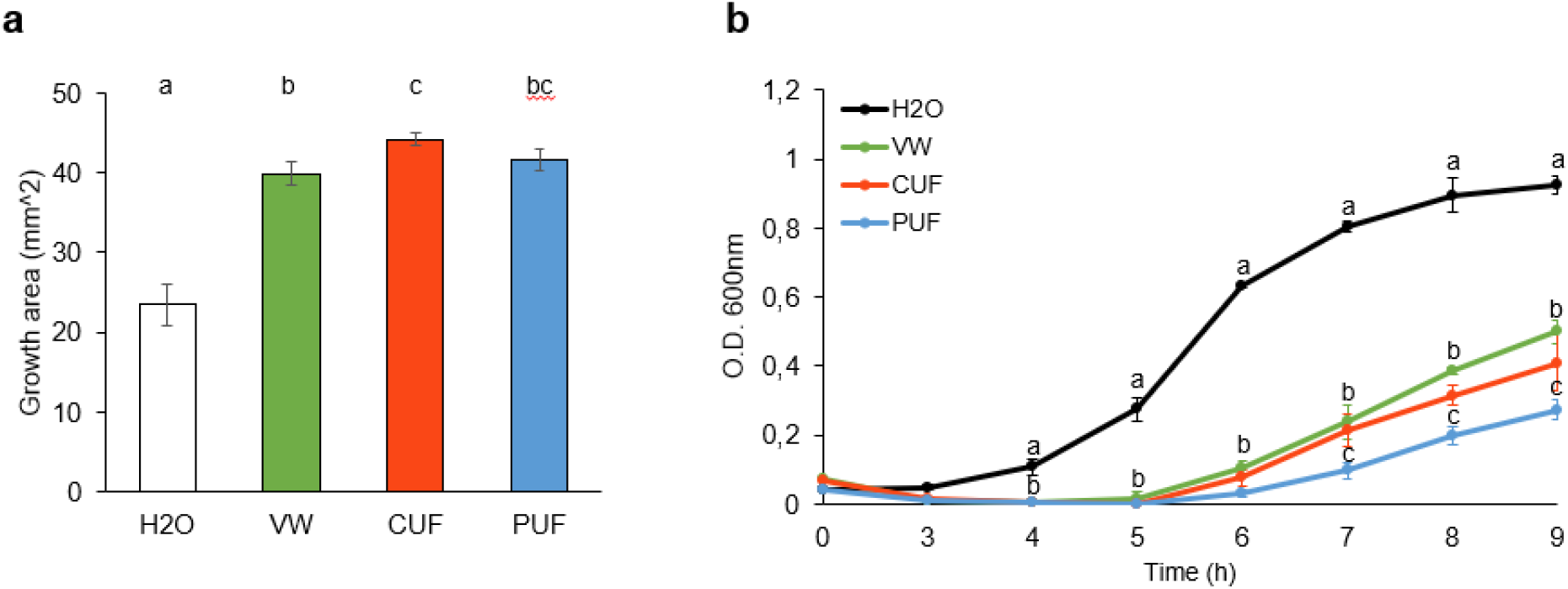
Effect of VW and TFMF fractions on *B. cinerea* and *P.carotovorum* growth in culture media. *B. cinerea* (a) and *P. carotovorum* (b) were grown respectively in PDB and LB medium supplied with 3 mg/ml of VW or TFMF fractions. The fungal growth area was measured after 48 h of incubation. These experiments were repeated three times with similar results. The data show mean ± SD (n=3). Different letters indicate P<0.05, as determined by ANOVA with Tukey’s HSD test.

Taken together, our results suggest the potential valorization of the VW-derived fractions as phyto-protectants in agriculture.

### 3.8. Treatment of *A. thaliana* plants with VW, CUF and PUF does not affect the vegetative growth and the plant fitness

One of the most noticeable physiological consequences of the activation of defense mechanisms in plants is the growth inhibition, according to the phenomenon known as growth defense trade-off (Huot *et al*., 2014) that is thought to occur in plants due to limited pool of resources that plant can invest either in growth or in defense to optimize fitness. In agricultural settings, crops have been bred for centuries to maximize growth-related traits resulting in a loss of genetic diversity that often compromises defense resulting in huge crop and economic losses (Strange and Scott, 2005). The modern need to meet rising global food demand requires to optimize the growth–defense balance to maximize crop yield. Thus, we evaluated the effect of VW and its TFMF derived fractions of interest on vegetative growth and plant fitness in Arabidopsis. VW, CUF and PUF, sprayed on adult plants at the concentration that induce pathogen resistance, did not affect plant growth (fig.8A) and fitness (fig. 8B and Table S4).

**Figure 8.**
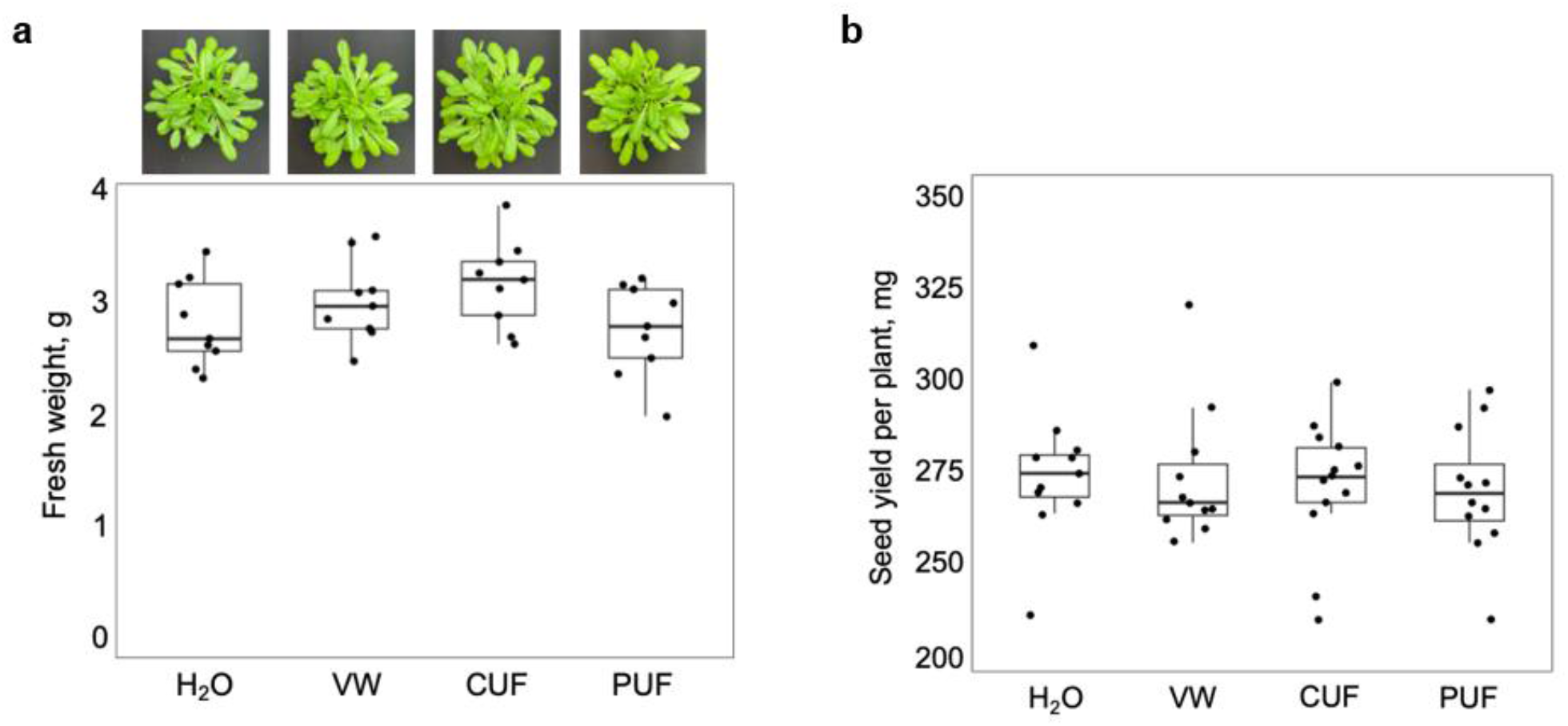
Treatment with VW and its TFMF derived fractions does not affect Arabidopsis vegetative growth and fitness. Four-week-old Arabidopsis plants were treated spraying VW, CUF and PUF at a concentration of 3 mg/mL. Plants treated with H_2_O were used as control. The fresh shoot weight per plant determined two weeks after treatment (a) and the seed yield per plant were evaluated (b). The data show mean ± SD (n=3). No statistically significant differences were found among treatments according to ANOVA with Tukey’s HSD test.

## 4. CONCLUSION

The olive oil industry generates substantial quantities of waste products, both solid and liquid, within a short timeframe of four months. Liquid waste, particularly VW, exhibits high pollutant and phytotoxic levels, posing significant challenges to the sector due to disposal difficulties and associated costs. The last generation Multiphase decanter technology with its novel Pâté by-product is now emerging in the olive oil extraction processes for the production of high-quality virgin olive oil (Lanza *et al*., 2020, Foti *et al*., 2021). Pâté without traces of stones, containing lipophilic and hydrophilic fractions is enriched in bioactive molecules, including anthocyanins, flavonoids, phenolic compounds and oligosaccharides with interesting functional properties and numerous potential industrial applications (Foti *et al*., 2021, Cuffaro *et al*., 2023, Shabir *et al*., 2023).

By recovering vegetation water after centrifugation and applying sequential stages of "solvent-free" TFMF procedure to VW, we successfully separated fractions enriched in bioactive molecules based on their dimensional and/or chemical-physical properties. In particular we found, in addition to bioactive phenols such as hydroxytyrosol, tyrosol and oleuropein, bioactive pectic fragments that had never been previously highlighted in the olive VW. OGs with DP comprised between 10 and 17 are well known elicitors of immune responses in plants against biotic stresses (Rakoczy-Lelek *et al*., 2023, Bigini *et al*., 2024, Radkowski *et al*., 2024). It was also reported that untreated OMW irrigation or fertilization can have toxic effects on plant growth, presumably due to high salinity, low pH and very high concentration of phenolics and fatty acids (Shabir *et al*., 2023). The absence of detrimental effects of TFMF selected fractions observed on plant growth and fitness and their positive effects in inducing immunity against pathogens, suggests their possible exploitation in commercial formulations as natural phyto-protectants sprays.

Notably, OGs have been included of the natural formulation of commercial plant protection products recognized by the EU indicating OG as “the first low-risk active substance” according with Regulation (EC) No 1107/2009 by the European Commission (EUR-Lex.europa.eu, 2015). In conclusion we have demonstrated, for the first time, that specific Pâté-derived fractions enriched in bioactive OGs induce [Ca^2+^]_cyt_ transients and defense gene expression and improve resistance against microbial pathogens in Arabidopsis model plants. We pave the way for field experiments with plants of agronomic interest for exploitation of these by-products as sustainable natural phyto-protectants in the control of pathogens causing devasting diseases of economically important crops (Dean *et al*., 2012, Davidsson *et al*., 2013) according to a circular economy perspective in agriculture.

## Funding statement

This work was supported by LazioInnova ("Progetti di Gruppi di Ricerca LR13/2008" ABASA (Agricultural By-products into valuable Assets for Sustainable Agriculture) funded to D.B., Sapienza University of Rome “Medi_Progetti_2023 (RM123188F796E238) funded to D.P., and Sapienza “Progetti per Avvio alla Ricerca” (AR1231888D22C2F6) funded to A.S.

## CRediT authorship contribution statement

**Ascenzo Salvati:** Writing – review & editing, Writing – original draft, Data curation, Investigation, Formal analysis, Software. **Fabio Sciubba:** Writing – review & editing, Investigation, Formal analysis, Software. **Alessandra Diomaiuti:** Investigation, Formal analysis. **Gian Paolo Leone** Investigation, Formal analysis. **Daniele Pizzichini:** Resources, Investigation, Formal analysis, Writing – original draft. **Daniela Bellincampi:** Resources, Conceptualization, Data curation, Writing – review & editing, Writing – original draft, Funding acquisition. **Daniela Pontiggia:** Conceptualization, Methodology, Data curation, Visualization, Investigation, Formal analysis, Funding acquisition, Supervision, Writing – original draft, Writing – review & editing.

## Data availability

The authors confirm that the data Sup. the findings of this study are available in the article and its supplementary materials.

## Declaration of competing interest

The authors declare that they have no known competing financial interests or personal relationships that could have appeared to influence the work reported in this paper.

## Acknowledgments

We also acknowledge the Sapienza Research Infrastructure NMR-based Metabolomics Laboratory. We are grateful to Dr. Barbara Bartolacci for addressing olive maturity index, for agronomic support and relations with local farms.

*A. thaliana* seeds stably expressing the Ca^2+^ biosensor GCaMP3 were a kind gift of Prof. Alex Costa of the University of Milan. We thank Prof. Giulia De Lorenzo of the Sapienza (University of Rome) for the support and helpful discussions.

## Supplementary materials

**Supplementary Table 1. Primers used in this work.**

**Supplementary Table 2: NMR Resonance assignment**

**Supplementary Table 3. Active OGs amount in PMF and CUF.** Data show the mean from three biological replicates ± standard deviation. DP; degree of polymerization

**Supplementary Table 4. Arabidopsis seeds dimensional parameters.** Data show the mean from minimum 12 biological replicates ± standard deviation. Two independent experiments with 6 replicates each were pooled. Not statistically significant differences were found among treatments.

**Supplementary Figure S1 HPAEC-PAD analysis of VW and different TFMF fractions compared with purified OGs mixture**. Signal intensities (nC) at each retention time (min) are indicated; standard oligomers are also shown with degree of polymerization (DP). VW, vegetation water; CMF, microfiltration concentrate; PUF, ultrafiltration permeate.

## Abbreviations

CW: cell wall
CMF: concentrate of microfiltration
CUF: concentrate of ultrafiltration
PMF: permeate of microfiltration
PUF: permeate of ultrafiltration
DMF: Multi-Phase Decanter
HPAEC-PAD: High-Performance Anion Exchange Chromatography with Pulsed Amperometric Detection
MF: microfiltration
VCR: volume concentratio ratio
NMR: nuclear magnetic resonance
OGs: oligogalacturonides
OMW: olive mill wastewater
TFMF: Tangential-flow membrane filtration
UF: ultrafiltration
VW: Vegetation water

## Notes

### Competing Interest Statement

The authors have declared no competing interest.

